# Stable isotope informed genome-resolved metagenomics reveals that Saccharibacteria utilize microbially processed plant derived carbon

**DOI:** 10.1101/211649

**Authors:** Evan P. Starr, Shengjing Shi, Steven J. Blazewicz, Alexander J. Probst, Donald J. Herman, Mary A. Firestone, Jillian F. Banfield

## Abstract

**Background:** The transformation of plant photosynthate into soil organic carbon and its recycling to CO_2_ by soil microorganisms is one of the central components of the terrestrial carbon cycle. There are currently large knowledge gaps related to which soil-associated microorganisms take up plant carbon in the rhizosphere and the fate of that carbon.

**Results:** We conducted an experiment in which common wild oats (*Avena fatua)* were grown in a ^13^CO_2_ atmosphere and the rhizosphere and non-rhizosphere soil was sampled for genomic analyses. Density gradient centrifugation of DNA extracted from soil samples enabled distinction of microbes that did and did not incorporate the ^13^C into their DNA. A 1.45 Mbp genome of a Saccharibacteria (TM7) was identified and, despite the microbial complexity of rhizosphere soil, curated to completion. The genome lacks many biosynthetic pathways, including genes required to synthesize DNA *de novo*. Rather, it requires externally-derived nucleotides for DNA and RNA synthesis. Given this, we conclude that rhizosphere-associated Saccharibacteria recycle DNA from bacteria that live off plant exudates and/or phage that acquired ^13^C because they preyed upon these bacteria and/or directly from the labelled plant DNA. Isotopic labeling indicates that the population was replicating during the six-week period of plant growth. Interestingly, the genome is ~30% larger than other complete Saccharibacteria genomes from non-soil environments, largely due to more genes for complex carbon utilization and amino acid metabolism. Given the ability to degrade cellulose, hemicellulose, pectin, starch and 1,3-β-glucan, we predict that this Saccharibacteria generates energy by fermentation of soil necromass and plant root exudates to acetate or lactate. The genome encodes a linear electron transport chain featuring a terminal oxidase, suggesting that this Saccharibacteria may respire aerobically. The genome encodes a hydrolase that could breakdown salicylic acid, a plant defense signaling molecule, and genes to make a variety of isoprenoids, including the plant hormone zeatin.

**Conclusions:** Rhizosphere Saccharibacteria likely depend on other bacteria for basic cellular building blocks. We propose that isotopically labeled CO_2_ is incorporated into plant-derived carbon and then into the DNA of rhizosphere organisms capable of nucleotide synthesis, and the nucleotides are recycled into Saccharibacterial genomes.

## Background

The Candidate Phyla Radiation (CPR) comprises a large fraction of the bacterial domain[1, 2]. An episymbiotic lifestyle has been demonstrated experimentally for only one group within the CPR, a member of the phyla Saccharibacteria, formerly TM7 [3]. Bacteria from this phylum are detected across a diversity of environments, suggesting that they may vary in their metabolic capacities. Originally, discovered in a peat bog [4], these bacteria have been described from human mouths [3], aquifers [5], sediment [6], and soil [7]. 16S rRNA gene surveys indicate that Saccharibacteria also occur in the rhizosphere, but little is known about their metabolism and how they differ from related organisms growing in other environments [8, 9].

There are four previously reported, closed and circularized genomes for Saccharibacteria (from a sludge bioreactor [10], human mouth [3] and two from sediment [11]) and some partial genomes. Based on genomic analyses, it was previously predicted that Saccharibacteria are anaerobic fermenters, despite several genomes encoding a cytochrome-o-ubiquinol oxidase (Complex IV). Complex IV is the final step in the aerobic electron transport chain, however, it has been suggested that Saccharibacteria use it for O_2_ scavenging, given the lack of other electron transport chain components. However, it has been documented that Saccharibacteria are able to grow in the presence of oxygen [12]. Here, we found that a Saccharibacteria population in rhizosphere soil incorporated ^13^C into its genomic DNA, indicating that this population was active and dividing. We report the complete genome for this organism and show how analysis of its metabolism sheds light on the pathway by which the label was incorporated. We compare the genome to those of other complete Saccharibacteria to address the question of how this bacterium is adapted to live in soil and specifically in the rhizosphere.

## Results

DNA was extracted from soil removed from the roots of an *Avena fatua* plant grown for six weeks in a ^13^CO_2_ atmosphere and separated into three fractions based on density. The isotopically light, middle and heavy fractions were sequenced and subjected to genome-resolved metagenomic analyses. From the rhizosphere middle fraction, accounting for 210 Mbp of contiguous sequence, a large scaffold was assembled *de novo* and could be circularized. Local assembly errors were identified and corrected and three scaffolding gaps were filled by manual curation. The complete, closed genome is 1.45 Mb in length with a GC content of 49.95%. The genome was most abundant in the middle fraction at 15x coverage, but was also present in the isotopically heavy and light fractions at ~3x coverage. The genome was undetectable in the background soil (**Figure 1**). DNA density is a function of both the extent of isotopic enrichment and the GC content, low GC is less dense than higher GC DNA [13]. This genome has a lower GC content than the rest of the assembly (average GC content of scaffolds larger than 1000 bp is 66%), which indicates ^13^C was incorporated, as nucleotides, into the DNA. The middle fraction where the genome was mainly detected had a density of 1.737-1.747 g/ml. Given that natural abundance DNA with 49.95% GC content would have a density of ~1.71 g/ml [14], we predict that the genome is at least 50% labeled.

**Figure 1.**
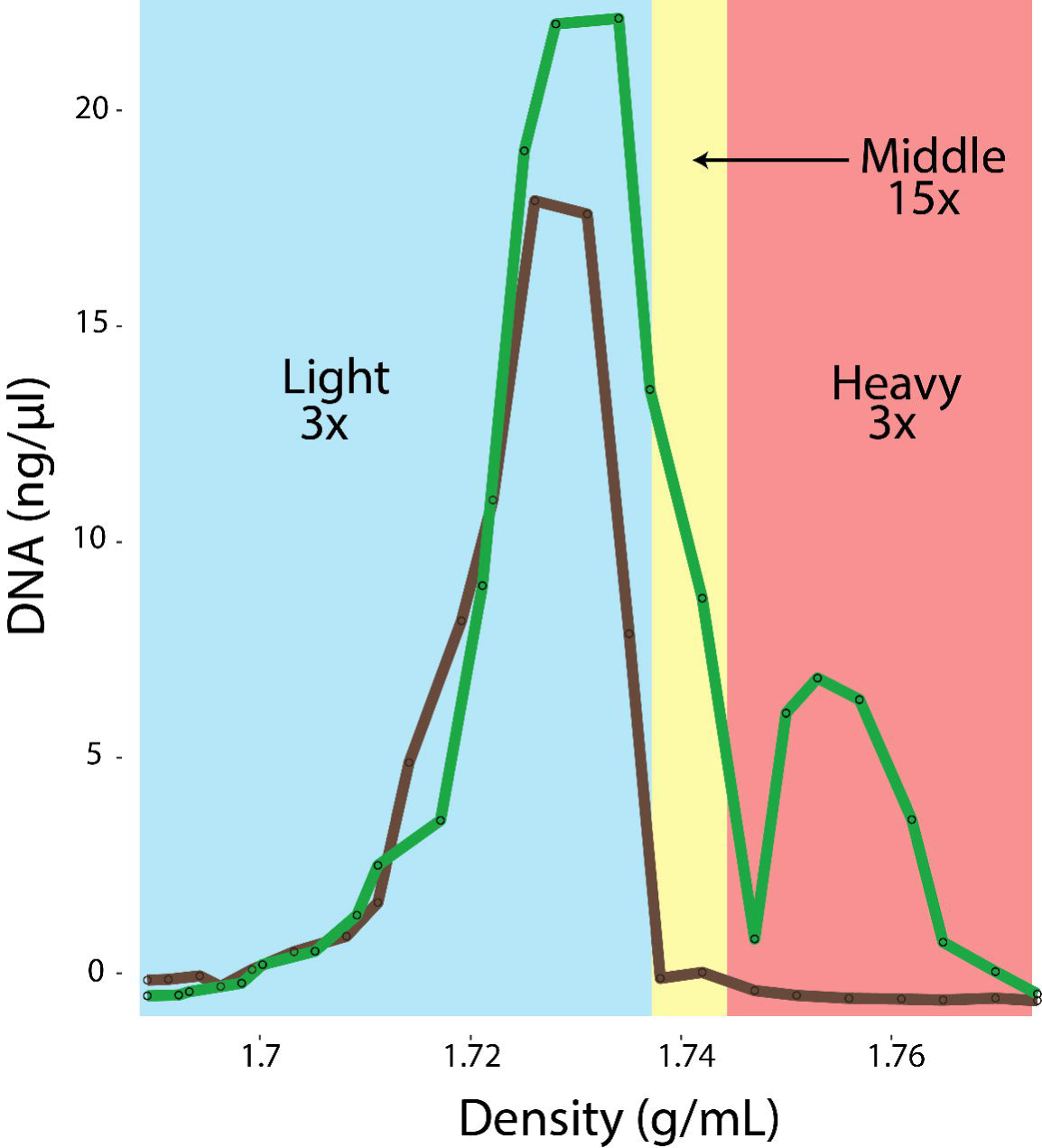
Stable isotope fraction determination. This plot shows the distribution of densities and concentrations of DNA extracted from rhizosphere and bulk soil following density centrifugation. The offset of the curves demonstrates clear enrichment of heavy DNA in the rhizosphere. The black dots on the curves represent individual fraction measurements. The three fractions are designated as light (blue shading), middle (yellow shading), and heavy (red shading). Numbers indicate the coverage of the *T. rhizospherense* genome in each fraction. The *T. rhizospherense* genome had < 1x coverage in each bulk soil fraction.

The genome has 1531 protein coding sequences (**Table S1**) and a full complement of tRNAs (46 in total). The 5S rRNA, 23S rRNA and 16S rRNA genes are in a single locus that also includes Ala and Ile tRNA genes. Based on the sequence of the 16S rRNA gene, the genome was assigned to be a member of the Saccharibacteria phyla. The closest 16S rRNA gene sequence in NCBI is from an unpublished study entitled “Effects of pinewood nematode (*Bursaphelenchus xylophilus*) infected-*Pinus massoniana* on soil bacterial communities”. The cloned 16S rRNA gene shares 94% identity with the sequence reported here (**Figure 2**). The most closely related genomically described organism is *Saccharimonas aalborgensis* from activated sludge with 84% identity across the full-length 16S rRNA gene. We propose the name *Candidatus* “Teamsevenus rhizospherense” for the organism described here, given the derivation of the genome from the rhizosphere. As this is the first genome from a distinct Class, the representatives of which mostly are found in soil, we propose the complete taxonomic descriptor: Phylum: *Candidatus* Saccharibacteria, Class: *Candidatus* Soliteamseven, Order: *Candidatus* Teamsevenales, Family: *Candidatus* Teamsevenaceae, Genus: *Candidatus* Teamsevenus, Species: *Candidatus* rhizospherense.

**Figure 2.**
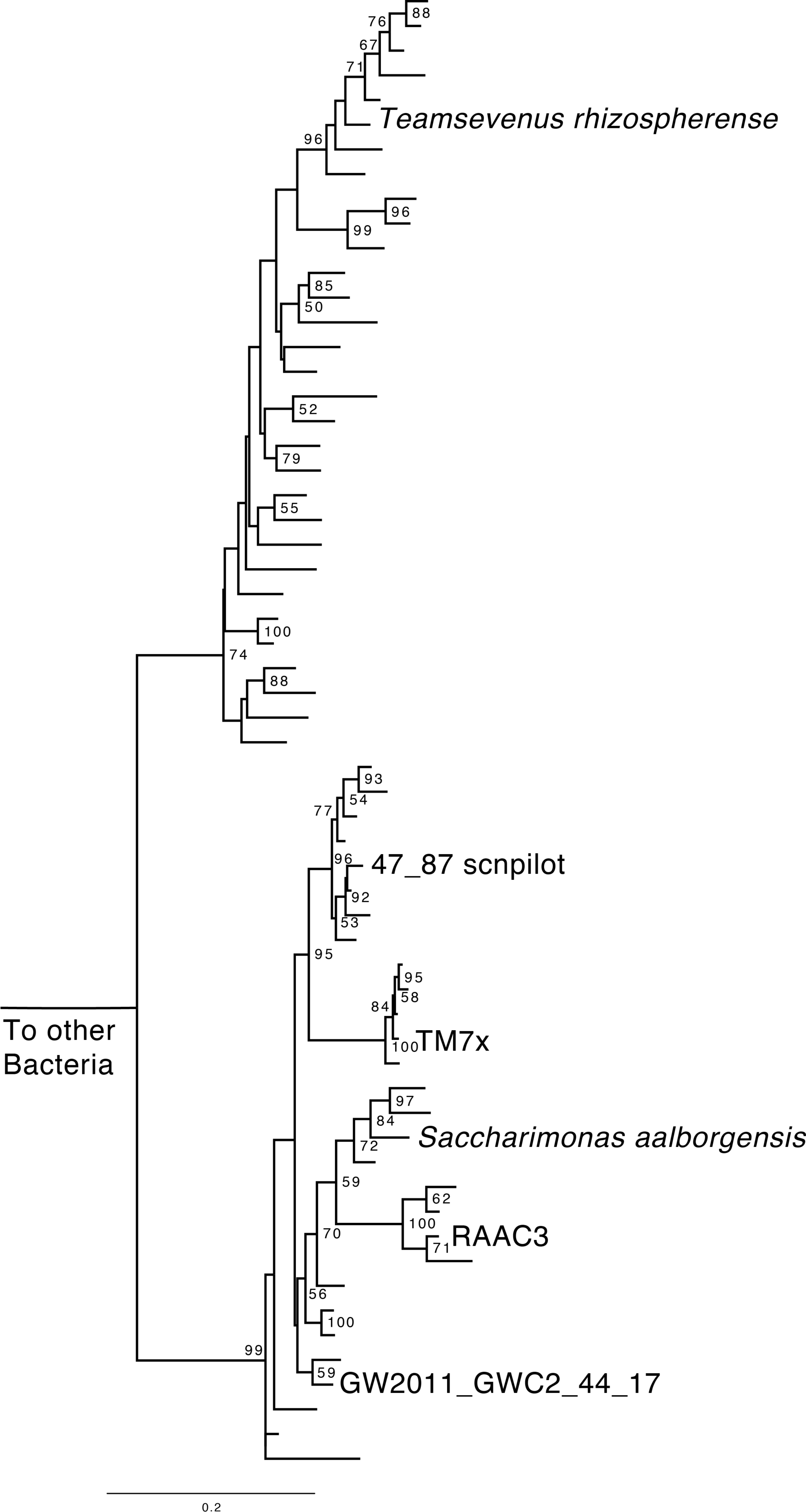
Phylogeny of Saccharibacteria based on 16S rRNA gene sequences. The maximum-likelihood tree shown was constructed from an alignment containing representative Saccharibacteria. Named branches indicate the complete genomes included in this study. Bootstrap values of >50% are displayed.

We calculated the GC skew and cumulative GC skew across the closed *T. rhizospherense* genome and found the symmetrical pattern typical for bacteria, with a single peak and trough indicative of the terminus and origin of replication (**Figure 3**). This result both validates the accuracy of the circularized genome and confirms that Saccharibacteria use the typical bacterial pattern of bi-directional replication from a single origin to the terminus (as do some Peregrinibacteria, another group of CPR bacteria[15]). The start of the genome was adjusted to correspond to the predicted origin, which lies between the DNA polymerase III subunit beta and the chromosomal replication initiator protein. It has a full set of ribosomal proteins, except for L30, which is uniformly absent in CPR bacteria [16].

**Figure 3.**
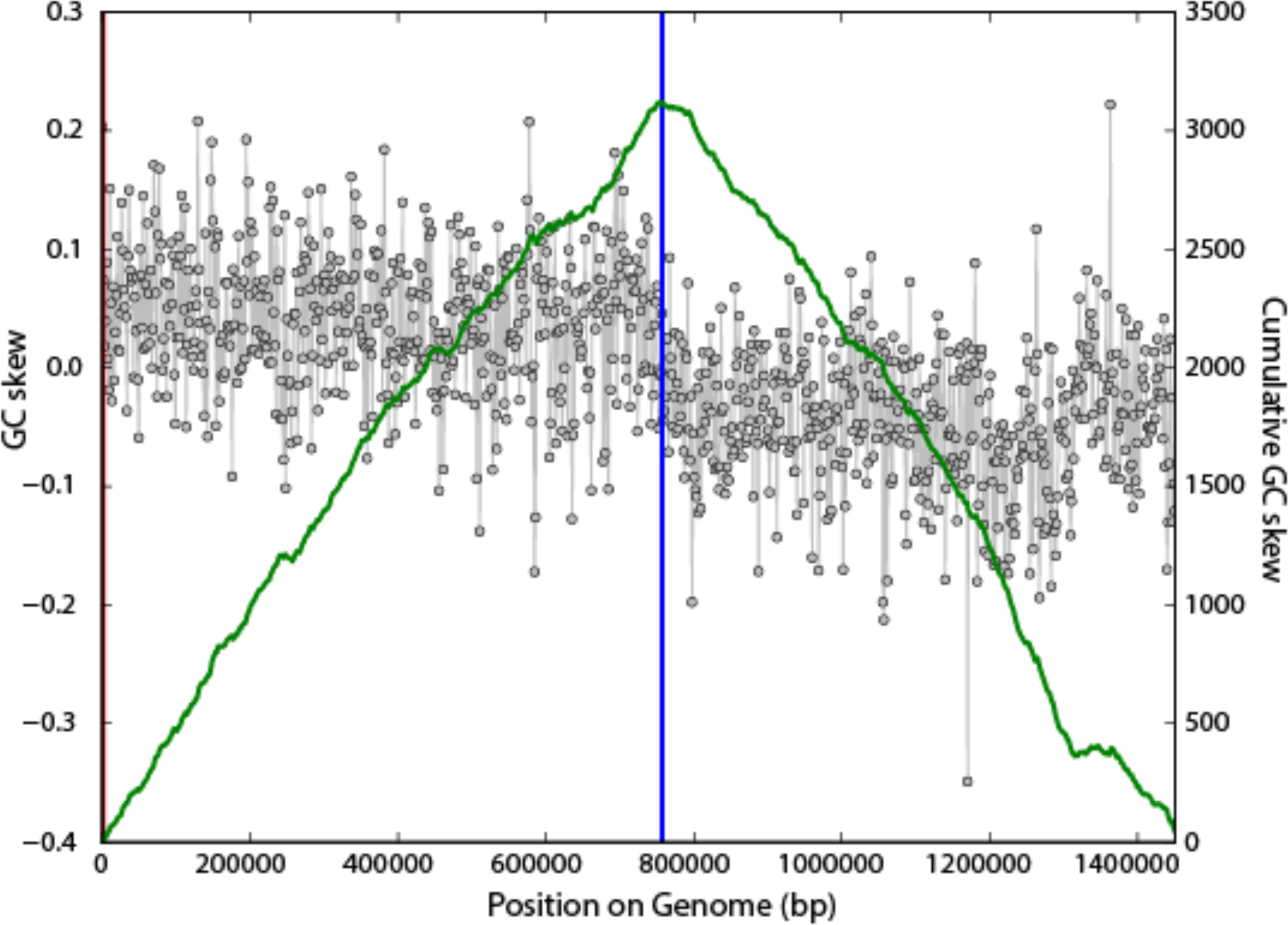
Plot of GC skew (black) and cumulative GC skew (green, window 1000 bp, slide of 10 bp) of the *T. rhizospherense* genome. The plot shows the predicted locations of the origin (red line, 1,201 bp) and terminus (blue line, 757,370 bp) of replication. The form of the plot is as expected for a correctly assembled, circularized genome that undergoes bi-directional replication from a single origin.

## Biosynthetic pathways

The *T. rhizospherense* genome encodes a number of enzymes for the conversion of nucleotides to NMP, NDP and NTP and formation of RNA. In addition, we identified genes to phosphorylate G, C and U. However, the organism seems unable to synthesize 5-phospho-alpha-D-ribose-1-diphosphate (PRPP). Further, it lacks essentially all of the steps for synthesis of nucleotide bases and the pathways that would convert PRPP to inosine monophosphate or uridine monophosphate. *S. rhizophera* may have a novel nucleotide biosynthesis pathway, but this is unlikely as nucleotide biosynthesis remains highly conserved across domains [17]. Thus, *S. rhizophera* likely did not *de novo* synthesize its nucleotides but rather must have acquired nucleotides from an external source. The genome encodes several nucleases, an external micrococcal nuclease, and an oligoribonuclease for the breakdown of externally-derived DNA and RNA (**Figure 4**). The mechanism for DNA and RNA import is unknown, as we did not identify nucleotide transporters. However, there are a number of transporters with unidentified specificity that could be involved in DNA or nucleotide uptake or the type IV pili could be responsible for this function. A large portion of the genome is dedicated to DNA and RNA repair mechanisms. There are twenty 8-oxo-dGTP-diphosphatase genes that prevent the incorporation of oxidized nucleotides. This may be an adaptation for life in a mostly aerobic environment, a seemingly unusual condition for members of the Saccharibacteria phylum, as all other complete genomes were found in mostly anaerobic environments and encode few genes with this function.

**Fig. 4.**
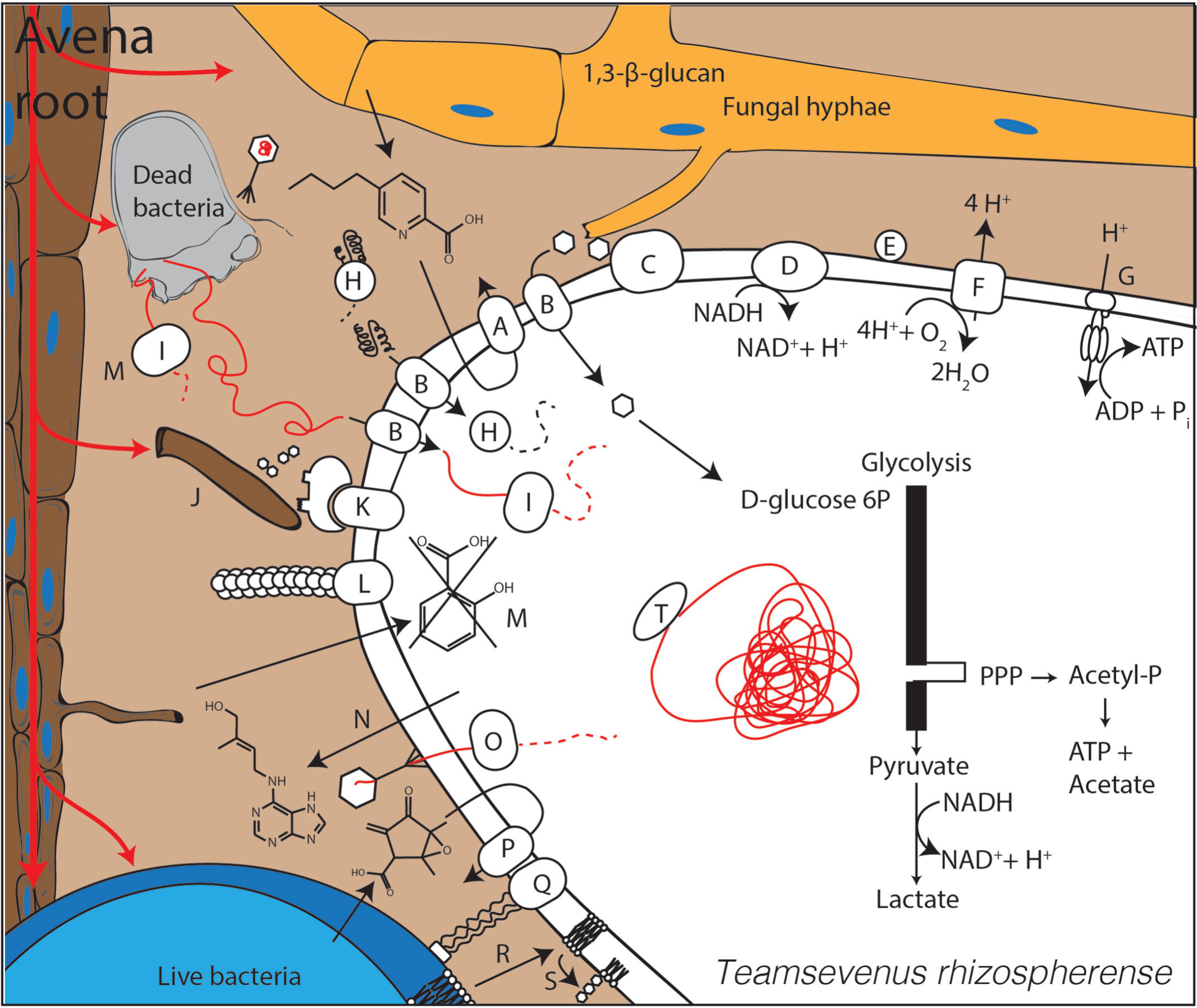
Cell diagram of *T. rhizospherense*. A. fusaric acid resistance machinery, B. unidentified importer, C. glucan 1,3-beta-glucosidase, D. NADH dehydrogenase II, E. blue-copper protein, F. cytochrome-o-ubiquinol oxidase, G. F-type H+-transporting ATPase, H. peptidase, I. nuclease, J. root hair, K. cellulosome, L. type IV pilus, M. salicylate hydroxylase, N. zeatin production, O. restriction system for possible phage defense, P. various antibiotic resistance mechanisms, Q. intercellular attachment, R. scavenging of lipids, S. production of phosphatidyl myo-inositol mannosides, T. DNA repair machinery.

The *T. rhizospherense* genome does not encode the ability to synthesize any amino acids *de novo* from central metabolites. However, it encodes genes to generate amino acids from precursors (e.g., valine and leucine from 2-oxoisovalerate, isoleucine from 2-methyl-2-oxopentanoate, and histidine from L-histidinol phosphate) and to interconvert some amino acids (serine and glycine). We identified genes for proteases that could breakdown externally-derived proteins. No amino acid specific transporters were annotated, but several transporters of unknown function could import the amino acids. There is little evidence to suggest that externally derived amino acids are broken down for use in the TCA (the only TCA cycle gene identified is a fumarate reductase subunit) or other cycles.

*T. rhizospherense* appears unable to synthesize fatty acids, yet it encodes three copies of the 3-oxoacyl-(acyl-carrier protein) reductase and five copies of acyl-CoA thioesterase I, indicating the capacity for fatty acid conversion. We predict that *T. rhizospherense* takes up phosphatidylcholine (possibly derived from eukaryotes), 1,2 diacyl sn-glycerol-3P (from bacteria), or phosphatidylethanolamine (the main bacterial phosopholipid) and may be able to interconvert the compounds using a gene annotated as phospholipase D. We identified a putative phosphatidate cytidylyltransferase that could add a head group to 1,2 diacyl sn-glycerol-3P forming CDP-diacylglycerol. This may be able to be converted into three products: phosphatidylglycerophosphate (via CDP-diacylglycerol-glycerol-3-phosphate 3-phosphatidyltransferase), or to cardiolipin (via cardiolipin synthase) or to phosphatidyl-1D-myoinositol (via CDP-diacylglycerol-inositol 3-phosphatidyltransferase).

Interestingly, phosphatidyl-1D-myo-inositol is the precursor for generation of phosphatidylinositol mannosides, glycolipids that are decorated by a chain of mannose molecules and that are found in the cell walls of *Mycobacterium* [18]. There are several genes for the first step in phosphatidylinositol mannosides biosynthesis, which involves modification of phosphatidyl-1D-myo-inositol by addition of mannose. These include phosphatidylinositol alpha-mannosyltransferase (three copies) and a single copy of alpha-1,6-mannosyltransferase. Subsequently, a polyprenol-P-mannose α-1,2-mannosyltransferase (CAZy glycosyltransferase family 87) adds another mannose group. Other mannose additions may involve the three copies of dolichol-phosphate mannosyltransferase, which transfer mannose from GDP-mannose to dolichol phosphate a mannose carrier involved in glycosylation. Thus, although *T. rhizospherense* appears to be unable to synthesize fatty acids, it appears that it may be able to generate membrane lipids, including phosphatidylinositol mannosides, if provided 1,2 diacyl sn-glycerol-3P.

The genome encodes a near-complete pathway for peptidoglycan biosynthesis. Given this, the one missing enzyme (murC, a ligase) may be too distant for identification and we predict the capacity for peptidoglycan synthesis.

This Saccharibacterial species appears to be reliant on external sources of cofactors. For instance, the genome encodes no genes in the nicotinate synthesis pathway, only one gene in the synthesis pathways for thiamine and riboflavin and two genes for the synthesis of biotin. However, there are a number of genes present for the biosynthesis of folate and the ability to activate, but not synthesize, B12 into its coenzyme form.

## Central metabolism and energy generation

Interestingly, the *T. rhizospherense* genome encodes a simple, two-subunit cellulosome that may be used to attach to and degrade plant or microbially-derived cellulose to cellobiose. Cellobiose is likely converted to D-glucose via one of 14 different glycosyl hydrolases. The genome also encodes genes for production of D-glucose via breakdown of starch/glycogen and trehalose. It also encodes several genes for hydrolysis of 1,3-β-glucan, one of the most common fungal cell wall polysaccharides [19], to D-glucose. Sugars are a significant portion of plant root exudation, these sugars could be fed directly into the *T. rhizospherense* metabolism [20]. D-glucose can be converted to D-glucose-6P and D-fructose-6P and fed into the glycolysis pathway. The genome lacks a gene to phosphorylate β-D-fructose-6P, but this reaction is accomplished via pentose phosphate pathway. The product of these reactions is pyruvate. Pyruvate is not converted to acetyl-CoA. Interestingly, no reaction appears to generate or use acetyl-CoA, as reported for Saccharibacteria RAAC3. Acetyl-CoA metabolism also appears to be lacking in the Saccharibacteria represented by the four other publically available complete genomes.

The *T. rhizospherense* genome encodes a phosphoketolase that converts D-xylose-5P derived from the pentose phosphate pathway to acetyl-P that then can be converted to acetate, with production of ATP. Thus, we predict growth via fermentation under anaerobic conditions. ATP also can be produced by a F-type H^+^ transporting ATPase.

In glycolysis, NAD^+^ is consumed to form NADH that can be regenerated via what appears to be a linear electron transport chain that includes a single subunit NADH dehydrogenase (NDH-II). We identified several residues expected for function. Modeling revealed a close secondary structure match between the *T. rhizospherense* protein and NDH-II characterized from *E. coli*. The large evolutionary distance between *T. rhizospherense* and *E. coli* likely accounts for differences in some active site residues.

The genome encodes a few genes in an incomplete pathway for production of quinone-based molecules, two of which are in multicopy (five copies of genes annotated as ubiG and two copies as ubiE). We suspect that quinone is scavenged from an external source. We identified genes for a cytochrome-o-ubiquinol oxidase. Subunit I contains nearly all functional residues as well as the residues that distinguish it from cytochrome-c oxidases [21]. Five FNR family transcriptional regulators may serve to detect O_2_. If O_2_ is available, electrons could be passed from the NDH-II to a complete cytochrome-o-ubiquinol oxidase, which pumps four protons with the reduction of oxygen.

Several genes such as a blue-copper protein and NADH-quinone oxidoreductase subunit L (NuoL)-related protein also may be involved in electron transfer and contribute to membrane potential in NDH-II systems [22]. Three cytosolic NAD(P)H:quinone oxidoreductase genes were identified. These may reduce semiquinone (SQ) formed under conditions of high O_2_ availability to prevent reaction of SQ with O_2_ to form oxygen radicals [23]. We identified a novel protein predicted to be associated with the cytoplasmic membrane with two of the domains found in subunits of the Na^+^-translocating NADH-quinone reductase and an FeS cluster domain. We suspect that this may function in the electron transport chain, converting NADH to NAD^+^. The subunit F-like domain may oxidize NADH and transfer electrons to the iron-sulfur domain and the subunit B-like domain could form the Na^+^ translocating channel [24].

## Other predicted capacities indicative of lifestyle

The genome lacks CRISPR-Cas defense. However, we identified a restriction modification system that could serve this function.

We predict the capacity for twitching motility, guided by one of four chemotaxis response regulators. In addition to DNA uptake, type IV pili could be used for attaching to other cells, root surfaces or solids. There are also several genes for pseudo-pili, and an autotransporter adhesin that may also be involved in cellular adhesion. Also, annotated were two CAZy carbohydrate-binding module family 44 genes for binding to cellulose and the capacity for biosynthesis of cellulose that could be used to attach to plant surfaces [25].

We identified genes encoding for laccase and pyranose 2-oxidase, which may be used for lignin breakdown or detoxification of phenolics, and genes for the detoxification of lignin byproducts, including 3-oxoadipate enol-lactonase and 4-carboxymuconolactone decarboxylase [26].

Interestingly, we identified genes whose roles may be to modulate plant physiology. For example, we identified genes for the production of cis-zeatin (a plant hormone) from isoprenoid precursors (but a pathway for formation of the precursor isopentenyl-PP was not present). The genome encodes a protein annotated as “Lonely Guy,” a cytokinin (a plant growth hormone) activating enzyme. We also found a gene encoding salicylate hydroxylase, which breaks down salicylic acid, a plant defense signaling molecule.

Also identified was N-acyl homoserine lactone hydrolase, a gene for quorum quenching of other soil microbes and a gene to form 3',5'-cyclic-AMP, which may be involved in intracellular signaling. We predict the presence of genes that confer resistance to bacterially produced antibiotics (beta-lactams, streptomycin, oleandomycin, methylenomycin A, vancomycin, and general macrolides). A gene annotated as phosphatidylglycerol lysyltransferase may produce lysylphosphatidylglycerol, a membrane lipid involved in cationic antimicrobial peptide resistance [27]. We also found a gene that may confer resistance to fusaric acid, an antibiotic made by a common fungal pathogen of grass [28]. This fungus was found to be growing in the rhizosphere (data not shown). Interestingly, there is a gene that encodes a 2,487 amino acid protein that contains domains found in insecticidal proteins, Yen-Tc toxin, rearrangement hotspot repeats, and a protective antigen domain. The protein is predicted to be secreted and may be a toxin. The genome contains at least 16 genes related to toxin/antitoxin systems which have been implicated in stress response [29].

## Comparative genomics

The *T. rhizospherense* genome is 28% larger than the largest reported complete Saccharibacteria genome, which is from an anaerobic bioreactor. It is 38% larger than the average size of all complete Saccharibacteria genomes (**Table 1**). By re-annotating and analyzing these previously reported complete genomes we found that the *T. rhizospherense* genome encodes nearly twice as many unannotated genes as the other Saccharibacteria.

**Table 1.**
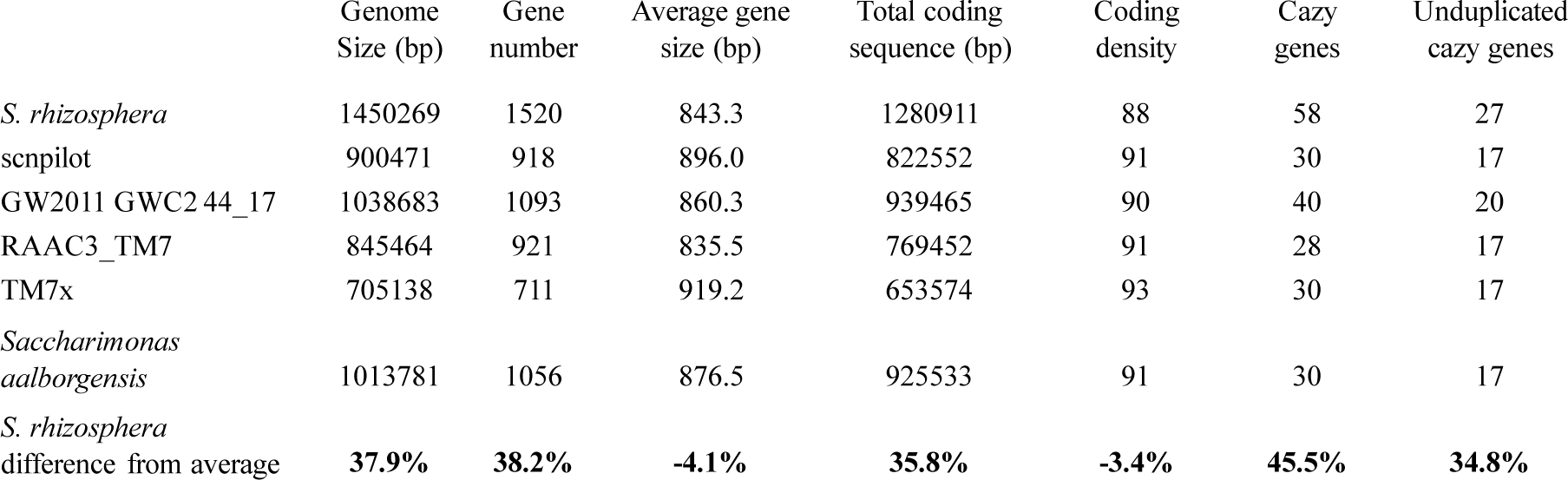
Genome statistics for the *T. rhizospherense* and other complete Saccharibacteria genomes

There are 130 *T. rhizospherense* functional annotations that were not found in any other Saccharibacteria, with 152 genes falling into these annotations. A few appear to be involved in amino acid metabolism, others in transcriptional regulation, DNA repair, sugar metabolism, transport, non-homologous end-joining, three genes annotated as NAD(P)H:quinone reductase and a dihydropteroate synthase for use in folate synthesis. The genome appears to encode a nickel superoxide dismutase for oxidative stress response. None of the other genomes encode the plant hormone related genes: “Lonely Guy” or salicylate hydroxylase. The genome also encodes transaldolase, an addition to the pentose phosphate pathway, pyruvate, water dikinase and the ability to take methylglyoxal to L-Lactate. It is the only Saccharibacteria with a twin-arginine protein translocation system (TatBC), although the function of the proteins with predicted TAT motifs are unknown. Of the *T. rhizospherense* genes involved in phosphatidylinositol mannoside production, CDP-diacylglycerol-inositol 3-phosphatidyltransferase is unique to *T. rhizospherense* and alpha-1,6-mannosyltransferase is also found in *Saccharimonas aalborgensis*.

*T. rhizospherense* has a notably larger repertoire of carbohydrate active enzymes than occurs in the other Saccharibacteria, including 58 genes with 27 unique annotations. Included in the set and not found in other Saccharibacteria genomes are genes predicted to confer the ability to hydrolyze hemicellulose (AG-oligosaccharides and mannooligosaccharides), amino sugars (galactosaminide), and pectin (oligogalacturonides).

The *T. rhizospherense* genome lacks several annotated genes found in all other Saccharibacteria. *T. rhizospherense* is unable to make (p)ppGpp, an alarmone that down regulates gene expression. Also lacking is dihydrolipoyl dehydrogenase, a subunit of pyruvate dehydrogenase (the function of which is unclear in these bacteria). Additionally not found are a recJ gene that encodes for a single stranded DNA exonuclease, a glutamine amidotransferase involved in cobyric acid synthase, the murC gene involved in peptidoglycan synthesis, and the PilW gene involved in pilus stability.

## Discussion

The *T. rhizospherense* genome does not encode the capacity to generate the ribose backbone of DNA or bases and thus it must acquire nucleotides from external sources. Importantly, its DNA is labeled with ^13^C that likely originated from ^13^CO_2_ fixed by the plant over the six-week study period. There are several sources of labeled DNA that might have been available to *T. rhizospherense*. We suspect that it obtained DNA from bacteria that lived off plant-derived carbon. However, some of the *T. rhizospherense* DNA might have been from phage that killed bacteria that grew on the ^13^C-labeled plant exudates. This is plausible because we identified labeled phage DNA (data not shown). A comprehensive analysis of the stable isotope probing experiment and overall community composition will be described in a separate publication. Alternatively, *T. rhizospherense* might acquire DNA from fungi or the plant, but we consider this less likely given the predicted obligate association between Saccharibacteria and other bacteria.

As Saccharibacteria depend on extracellular nucleosides, they may have difficulty regulating the correct nucleoside concentrations. This is known to be a mutagenic condition and a possible reason for the large number of DNA repair mechanisms [30]. The lifestyle of *T. rhizospherense* is novel compared to those of other CPR in that it is predicted to have the capacity to grow aerobically. Consistent with this, it has a large number of oxidative stress response genes. A close relationship with plant roots also may explain the large number of these genes, as plants release reactive oxygen species in response to pathogens and stress. It is the first member of the Saccharibacteria to be described from the rhizosphere and it appears to be adapted to life there via an expanded genetic repertoire for carbohydrate degradation relative to other Saccharibacteria.

It is possible that *T. rhizospherense* is a symbiont of a soil dwelling bacterium, with a lifestyle analogous to that of the Saccharibacteria TM7x that was cultured as an epibiont of a mouth dwelling Actinobacteria [3, 31]. A similar association is indicated by the antibiotic resistance mechanisms that are encoded by *T. rhizospherense* that protect against compounds produced by Actinobacteria, methylenomycin A, oleandomycin, vancomycin and streptomycin. Additionally, the ability for *T. rhizospherense* to produce phosphatidylinositol mannosides provides an intriguing connection between this organism and the clades of Actinobacteria that produce phosphatidylinositol mannosides, *Mycobacterium* being the best studied. In *Mycobacterium*, this cell wall component may protect the cell from antibiotics and act as a virulence factor during infection [18].

We do not know the extent of the labeling of cellular components of *T. rhizospherense,* other than the DNA, so we cannot rule out the possibility that this organism makes direct use of plant exudates for purposes other than DNA synthesis. However, the presence of machinery to degrade a variety of complex carbohydrates including cellulose and fungal cell walls suggests that it may not rely completely on plant exudates. *T. rhizospherense* has many genes for attachment that could be used to connect to a microbial host or plant surfaces. It may directly interact with the plant, given genes for the modulation of signaling molecules zeatin and salicylic acid.

## Conclusions

To our knowledge, this is the first study to generate a complete, circularized genome from a soil dwelling bacteria *de novo* from any soil metagenome. It is the first stable isotope-informed genome-resolved metagenomic study in the rhizosphere and thus the first soil microbiome study to make use of stable isotope probing to track carbon from atmospheric CO_2_ into the genome of a novel rhizosphere organism. The evidence indicates that ^13^C was incorporated into the *T. rhizospherense* DNA through its use of nucleic acids that were synthesized by other organisms that took up plant exudates. *T. rhizospherense* is the first genomically described rhizosphere-associated member of the Saccharibacteria phylum and many of its predicted metabolic capacities distinguish this organism from related Saccharibacteria that live in anaerobic environments. The rhizosphere alternates between aerobic and anaerobic conditions, and it appears that *T. rhizospherense* may have adapted to these cycles by having the capacity to perform fermentation and to grow aerobically using an unusual electron transport chain.

## Materials and methods

### Labelling

Soil (0-10cm) was collected from the University of California Hopland Research and Extension Center (Hopland, CA, USA), from an area where *Avena* spp. are a common grass. Microcosms were constructed and plant growth conditions were regulated as described previously [32]. The *A. fatua* plants were grown in labelling chambers with a 400 ppm atmosphere, the native CO_2_ was replenished with 99 atm% ^13^CO_2_. Rhizosphere and bulk soils were sampled at vegetative (6 weeks) and flowering (9 weeks) stages; microcosms were destructively harvested for rhizosphere soil and from bulk soil bags which exclude root ingrowth. Rhizosphere soil was washed off the roots and DNA was extracted from 0.5g of soil using a phenol:chloroform extraction detailed in [32].

### Stable isotope probing

To separate isotopically enriched DNA from unenriched DNA, each sample was separated based on density in a cesium chloride density gradient formed in an ultracentrifuge. Gradients were generated according to the method previously described [33]. Briefly 5.5 µg of DNA was added to the gradient buffer to create a solution with a density of 1.735 g/mL. Then 5.2 mL of the solution was transferred to an ultracentrifuge tube (Beckman Coulter Quick-Seal, 13 × 51 mm). Tubes were spun in an Optima L-90K ultracentrifuge (Beckman Coulter, Brea, California, USA) using a VTi65.2 rotor at 44000 rpm (176284 RCF_avg_) at 20°C for 109 h with maximum acceleration and braking. Then the content of the ultracentrifuge tube was separated into ~32 fractions using a syringe pump to deliver light mineral oil at 0.25mL/min to displace the gradient solution from the pierced bottom of the tube. Each fraction was approximately 12 drops (~144µL). The density of each fraction was measured using an AR200 digital refractometer (Reichert Inc., Depew, New York, USA). The DNA for each fraction was precipitated and quantified as previously described [33]. Fractions were then binned based on density and by comparison between the rhizosphere samples and the associated bulk sample (light= 1.692-1.737 g/mL; middle= 1.738-1.746 g/mL; heavy= 1.747-1.765 g/mL)

### Sequencing

Each fraction, light, middle, and heavy was then sent to the UC Davis Genome Center DNA Technologies Core for sequencing. Each sample was sequenced using an Illumina HiSeq 3000 (Illumina Inc., Hayward, California, USA) with paired-end libraries prepared with the Kapa Hyper protocol and a read length of 150 bp.

### Genome reconstruction, annotation and analysis

Reads were trimmed using Sickle (https://github.com/najoshi/sickle); BBtools (https://sourceforge.net/projects/bbmap/) was used to remove Illumina adapters and trace contaminants; finally reads were assembled using IDBA-UD (-step 20, -maxk 140, -mink 40) [34]. A single 1.45 Mb scaffold was recovered and was able to be circularized, then scaffolding errors and completeness were assessed as described in [16] and manually by mapping reads to the scaffold using Bowtie2 on default settings [35]. The scaffold was visualized in Geneious [36]. Genes were predicted using Prodigal [37]. Predicted ORFs were functionally described using a multidatabase search pipeline. Sequence similarity searches were performed with USEARCH [38] against UniRef100 [39], Uniprot [40], and the KEGG database [41]. Additional gene annotations were assigned using HMMs that were constructed based on KEGG Ortohologies [41]. All the proteins assigned to a KO were clustered using MCL [42] with inflation parameter (-I) of 1.1, based on global percent identity. Clusters were aligned using MAFFT v7 [43], and HMMs were constructed using the HMMER suite [44]. Carbohydrate-active enzymes were identified using dbCAN [45]. Domain level functional annotations were done using InterProScan [46]. tRNAs were predicted using tRNAScan-SE [47] and cellular localization was predicted using PSORTb with the gram-positive setting, which was demonstrated though previous imaging [48, 49]. The GC skew of the genome was calculated based on previously published tools [50]. Protein modelling was done with Swiss Model and NDH-2 modelling incorporated *Caldalkalibacillus thermarum* NDH-2 protein structure [51, 52]. Representative Saccaribacteria 16S rRNA sequences were obtained from NCBI and aligned using ssu-align [53] then a maximum-likelihood tree was constructed with RAxML by using the GTRCAT model with 100 bootstraps.

## Declarations

**Ethics approval and consent to participate:** Not applicable.

## Consent for publication

Not applicable.

## Availability of data and material

The genome will be made available on NCBI upon acceptance of this manuscript and via <u>http://ggkbase.berkeley.edu/tm7cure/organisms/342358</u>. The mapped read file can be also be downloaded from this site.

## Authors' contributions

SS and MF designed the labeling experiment. SS and DH carried out the labeling. ES and SB conducted the density centrifugation. ES and JB analyzed the data, with input from AP. ES and JB wrote the manuscript and all authors read and approved the final manuscript.

## Acknowledgements

This work was made possible by samples and expertise provided by personnel at the Hopland Research and Extension Center (Hopland, CA, USA). Sequencing was done at the UC Davis Genome Center and technical advice was provided by Lutz Froenicke. The HMM pipeline was developed by Dr. David Burstein and protein modeling assistance was provided by Dr. Cindy Castelle.

---

Supplementary Table 1: Genome summary, including HMM-based annotations.

